## Supplementary figure 1 for "MHC1-TIP enables single-tube multimodal immunopeptidome profiling and uncovers intratumoral heterogeneity in antigen presentation"

### Supplementary figures

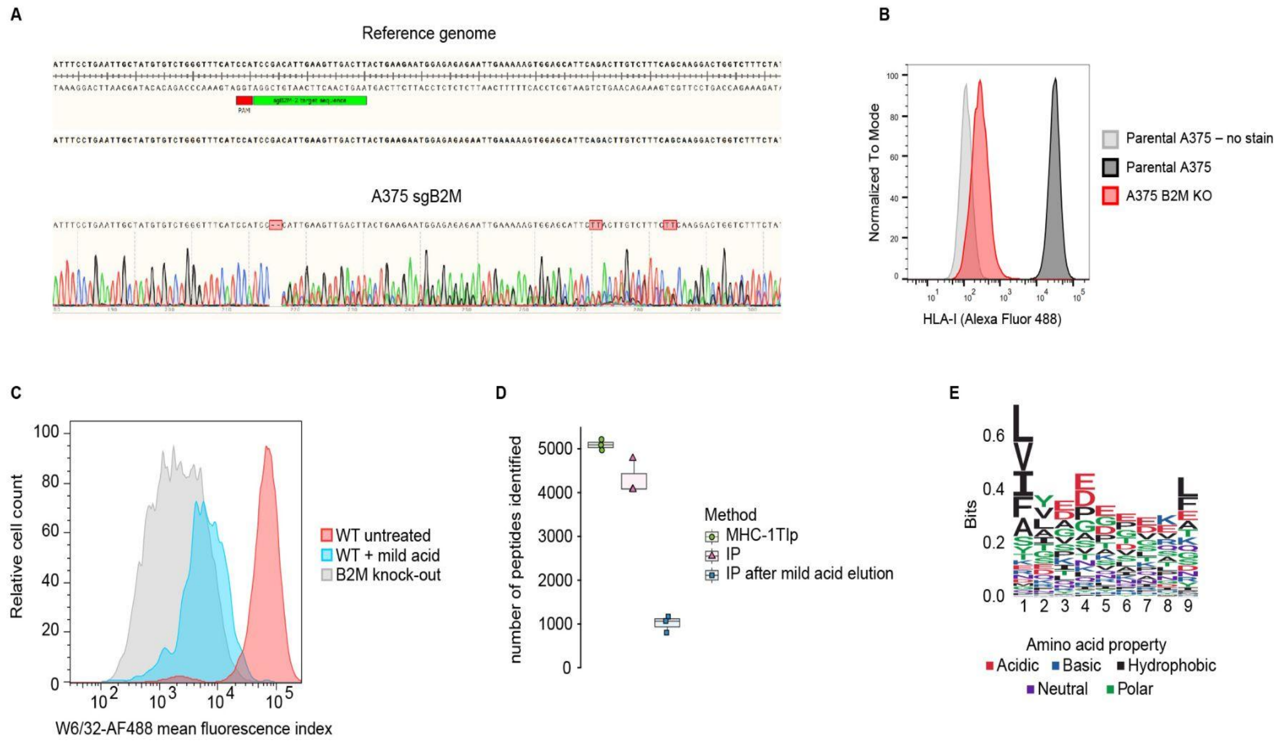


**Supplementary Figure 1: Recovery of intracellular immunopeptides and dispase-derived peptides.**

(A) Sanger sequencing Applied Biosystems Sequence Trace file alignment to sgB2M target sequence of the human reference genome hg38 in Snapgene v8.0.1.

(B) Flow cytometry staining with W6/32 antibody (pan-HLA-I)

(C) Flow cytometry staining with W6/32 antibody (pan-HLA-I)

(D) Number of immunopeptides identified with MHC1-TIP, immunoprecipitation (IP)-based immunopeptidomics and IP performed after mild acid elution in 10 million A375 cells

(E) Sequence motif generated from peptides of length 9, eluted from PDO-1 after dispase treatment to dissociate the organoids
