## Supplementary figure 2 for "MHC1-TIP enables single-tube multimodal immunopeptidome profiling and uncovers intratumoral heterogeneity in antigen presentation"

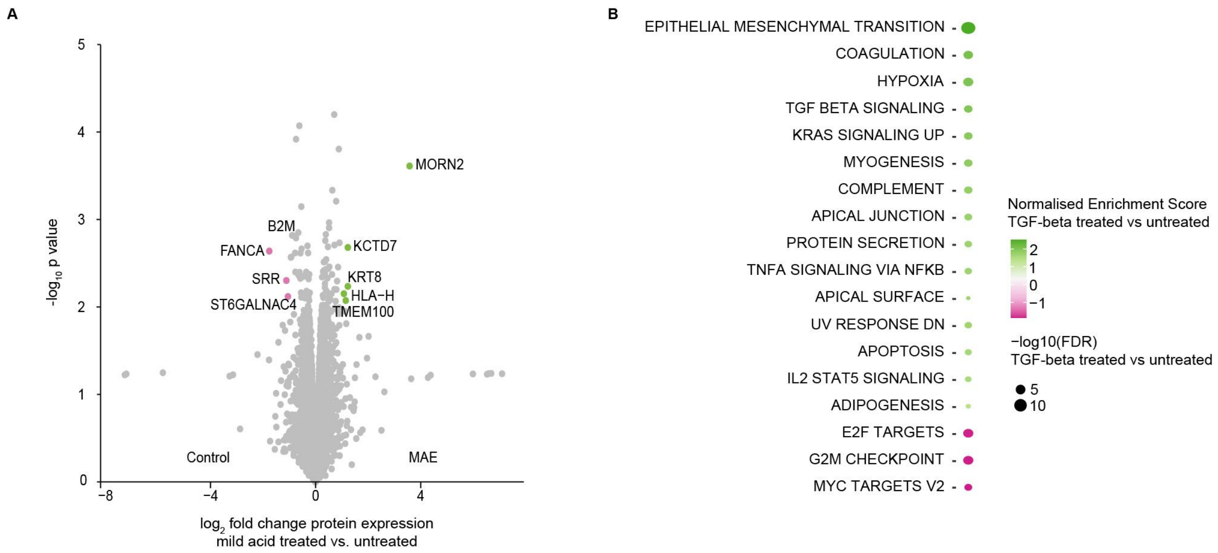


**Supplementary figure 2: MHC1-TIP enables multi-omic profiling**

(A) Changes induced in the proteome after mild acid elution. Green dots represent significantly upregulated proteins (FDR < 0.05 and $log_2$ fold change > 1) and pink dots represent significantly downregulated proteins (FDR < 0.05 and $log_2$ fold change < -1)

(B) Significantly enriched pathways (FDR < 0.05) after gene set enrichment analysis using Hallmark gene sets
