## Supplementary figure 3 for "MHC1-TIP enables single-tube multimodal immunopeptidome profiling and uncovers intratumoral heterogeneity in antigen presentation"

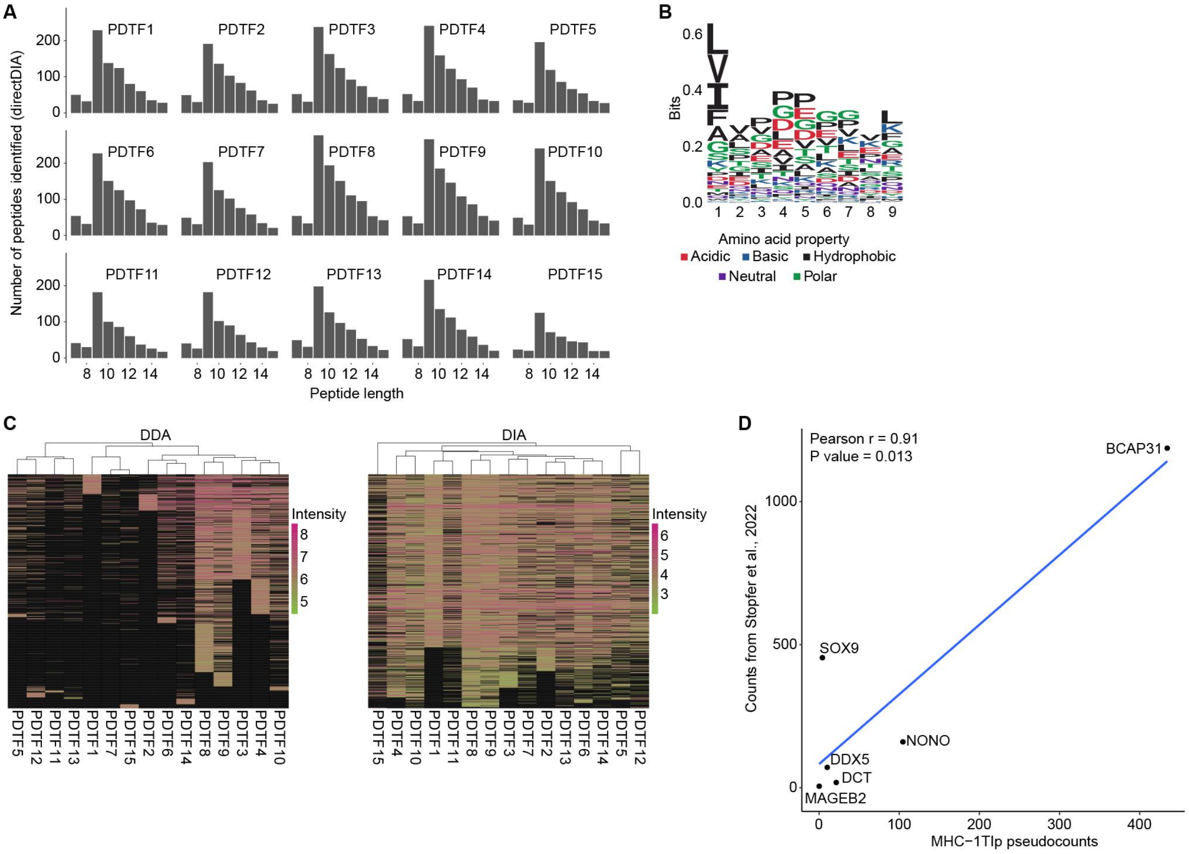


**Supplementary Figure 3: MHC1-TIP enables quantitative immunopeptidomics from tumour tissue fragments**

(A) Number of peptides detected per tissue fragment using direct DIA. All fragments show an enrichment of peptides of length 9

(B) Sequence motif of peptides of length 9 eluted from tissue fragments after dissociation of the fragment into a single-cell suspension using collagenase, DNAse and hyaluronidase

(C) Heatmaps showing data missing-ness (in black) with data-dependent and data-independent modes of acquisition from each tissue fragment. Intensities are plotted in $\log_{10}$ scale.

(D) Pseudocounts generated from MHC1-TIP data (immunopeptidomes of the untreated samples from the TGF-beta experiment; Figure 2) using a normalization method based on B2M protein copy numbers correlate with published hipMHC-quantified absolute antigen counts for the same cell type (A375) from Stopfer et al., 2022
