## Supplementary figure 4 for "MHC1-TIP enables single-tube multimodal immunopeptidome profiling and uncovers intratumoral heterogeneity in antigen presentation"

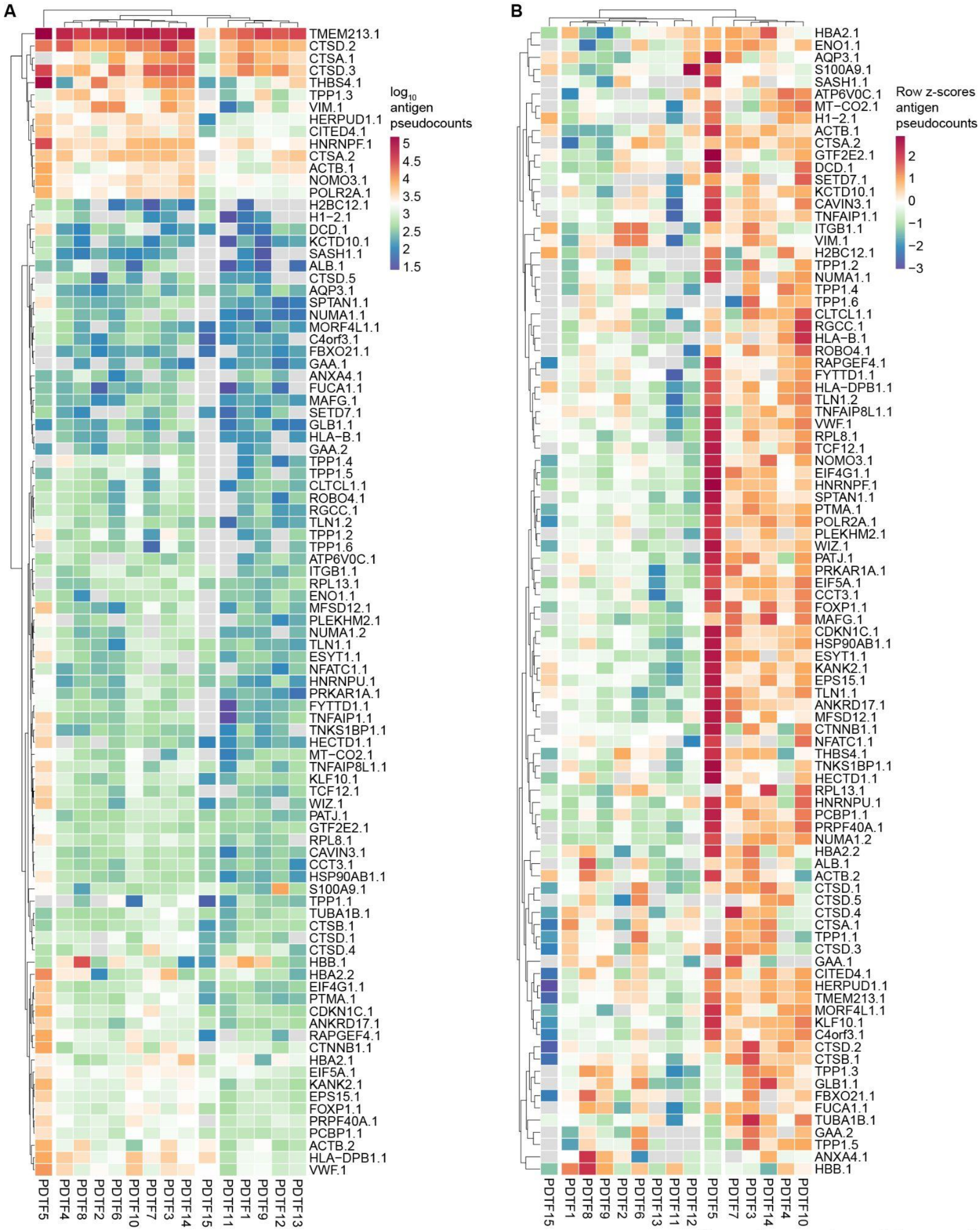


**Supplementary Figure 4: Patient-derived tumour fragments show heterogeneity antigen abundances.**

(A-B) Immunopeptides are labelled with the gene name of the parental protein of the antigen, conjugated with a numeric identifier for each peptide identified from that protein

(A) Heatmap displaying pseudocounts of antigen per cell in each tissue fragment. Detected antigen copy numbers per cell range 4 orders of magnitude.

(B) Scaling and centering antigen pseudocounts row-wise reveals heterogeneity in antigen presentation across the fragments
